# A negative feedback mechanism in the insulin-regulated glucose homeostasis in Japanese flounder *Paralichthys olivaceus* by two ways of glucose administration

**DOI:** 10.1101/173948

**Authors:** Dong Liu, Dongdong Han, Benyue Guo, Kangyu Deng, Zhixiang Gu, Mengxi Yang, Wei Xu, Wenbing Zhang, Kangsen Mai

## Abstract

The present study comparatively analyzed the blood glucose and insulin concentration, the temporal and spatial expression of brain-gut peptides and the key enzymes of glycolysis and gluconeogenesis in Japanese flounder by intraperitoneal (IP) injection and oral (OR) administration of glucose. Samples were collected at 0, 1, 3, 5, 7, 9, 12, 24 and 48h after IP and OR, respectively. Results showed that the hyperglycemia lasted 5 hours and 21 hours in OR and IP group, respectively. The serum insulin concentration significantly decreased (1.58±0.21mIU/L) at 3h after IP glucose. However, it significantly increased at 3h (3.37±0.34mIU/L) after OR glucose. The gene expressions of prosomatostatin, neuropeptide Y, cholecystokinin precursor and orexin precursor in the brain showed different profiles between the OR and IP group. The OR not IP administration of glucose had significant effects on the gene expressions of preprovasoactive intestinal peptide, pituitary adenylate cyclase activating polypeptide and gastrin in the intestine. When the blood glucose concentration peaked in both IP and OR group, the glucokinase expression in liver was stimulated, but the expression of fructose-1,6-bisphosphatase was depressed. In conclusion, brain-gut peptides were confirmed in the present study. And the serum insulin and the brain-gut peptides have different responses between the IP and OR administration of glucose. A negative feedback mechanism in the insulin-regulated glucose homeostasis was suggested in Japanese flounder. Furthermore, this regulation could be conducted by activating PI3k-Akt, and then lead to the pathway downstream changes in glycolysis and gluconeogenesis.

## 1. Introduction

Glucose acts as an important source of energy for most species of fish. However, carnivorous fish species like rainbow trout have been considered as “glucose intolerant” as hyperglycemia after a glucose load can last for several hours, even more than one day due to their limited ability in using glucose efficiently (Aguilar, Conde-Sieira et al. 2010). They rely more on lipid and protein for energy purposes. Fish has an oxygen consumption rate one-tenth that of mammals. It was also found that the ability of insulin secretion stimulating by dietary amino acids is stronger than that of glucose (Mommsen and Plisetskaya 1991). Meanwhile, the ability of glucose phosphorylation and glucose transport is weak in fish (Cowey and Walton 1989).

Although glucose is not the main energy substrate for carnivorous fish species, glucose metabolism is important for the function of specific tissues in fish, such as brain (Soengas and Aldegunde 2002). The brain has the highest glucose utilization rates per unit mass in all tissues examined in rainbow trout (Washburn, Bruss et al. 1992). In mammals, the brain is responsible for neuroregulation, meanwhile, gut cells perceive nutrients. The brain and gut have complicated connections in nutrients, hormone as well as nerve, thus a gut–brain axis has been generated (Romijn, Corssmit et al. 2008). Pearse and Takor (Pearse and Takor 1979) pointed out that gastrointestinal peptide secreting cells and peptidergic neurons in the brain is common in embryogenesis originated from the neuroectoderm. Some researches put forward the concept of enteric nervous system, and it has close ties to brain systems (Wood 1996). As the discovery of brain-gut peptides, researchers come up with the hypothesis of brain-intestinal connection (Zhang 2001). Although the function of the gut–brain axis is not completely understood yet in fish, the brain-peptides like gastrin (Pereira, Costa et al. 2015), cholecystokinin (Polakof, Míguez et al. 2011) and somatostatin (SS) (Sheridan, Plisetskaya et al. 1987) has been demonstrated to play a crucial role in glucose homeostasis.

Somatostatin promotes glycogen breakdown and the release of glucose from liver to plasma (Eilertson, O’connor et al. 1991). And it can also inhibit hormones secretion, including insulin (Sheridan, Plisetskaya et al. 1987, Eilertson and Sheridan 1993, Very, Knutson et al. 2001). Neuropeptide Y (NPY) is also involved in regulating insulin and glucose metabolism in fish. It was found that plasma insulin levels decreased in fasted European sea bass co-injected with NPY plus glucose, but remained stable when NPY was administrated alone to fed and fasted animals (Cerdá-Reverter, Sorbera et al. 1999). Mammalian studies showed that cholecystokinin (CCK) can stimulate insulin secretion and it can also make influence on β cell proliferation (May, Liu et al. 2016). The CCK promotes the transport of insulin into the central nervous system (CNS) of rats (Hermansen 1984). The Orexin (OX) is involved in glucose and lipid metabolism in mammals. Oerxin-A can promote insulin secretion, and it also enhances the capacity of glucose stimulated insulin (Park, Shim et al. 2015). In addition, the function of vasoactive intestinal peptide (VIP) and pituitary adenylate cyclase activating polypeptide (PACAP) are very similar. It mainly manifested in animal reproduction, nutrient digestion and energy balance. It deserves to be mentioned that these two hormones can stimulate insulin secretion (Bredkjoer, Palle et al. 1997, Vaudry, Gonzalez et al. 2000). Gastrin has the similar function to VIP/PACAP in glucose metabolism, and it can also stimulate insulin production (ShuangmingYue 2007). In regard to the glucose metabolism in mammalians, the relationship between the hormones mentioned above was reviewed in Figure 1. However, little related information was reported in fish.

**Figure 1.**
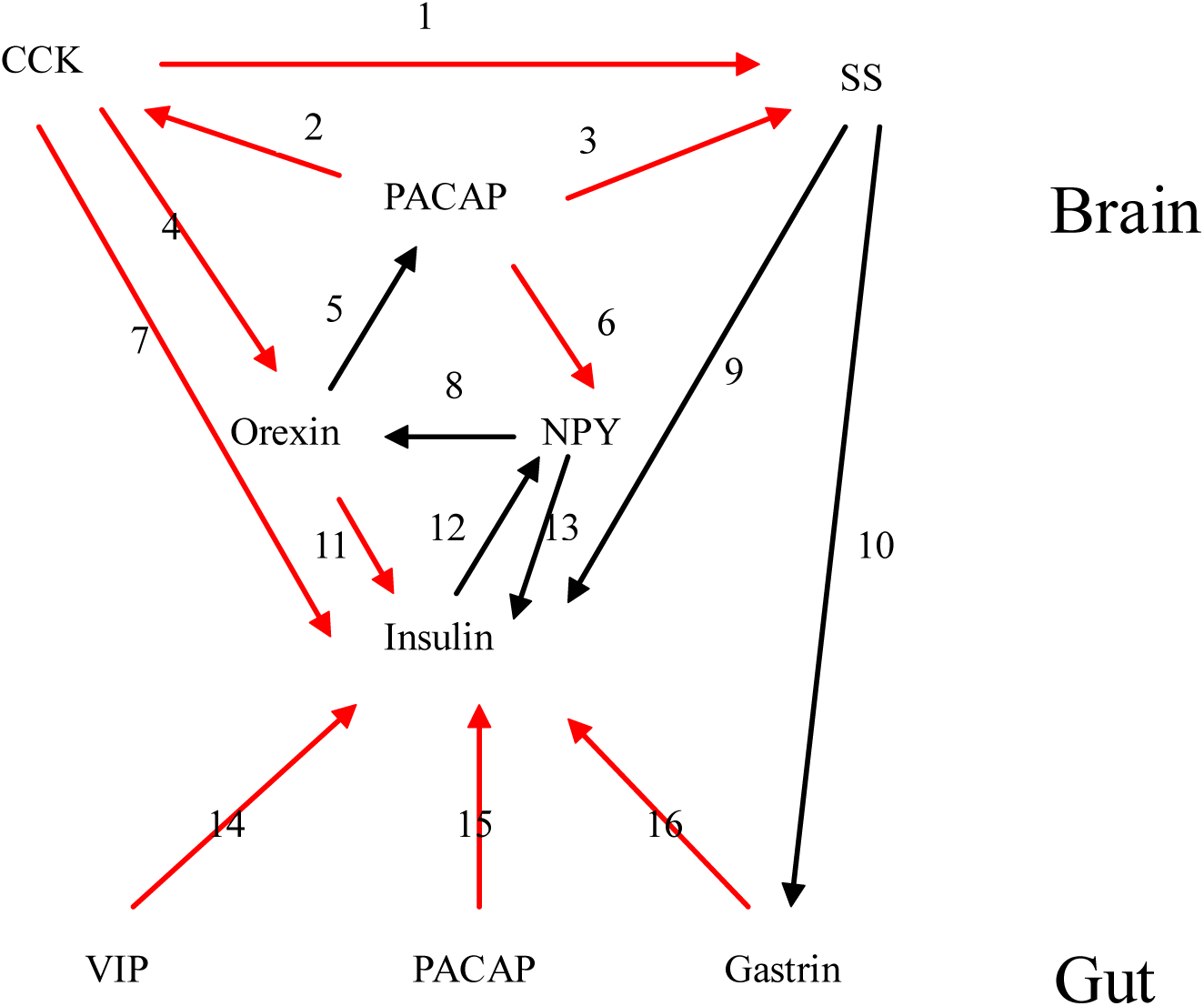
The relationship between different hormones in mammalians. Red line stands for promotion, and black line means inhibition. Somatostatin (SS), neuropeptide Y (NPY), cholecystokinin (CCK), vasoactive intestinal peptide (VIP), pituitary adenylate cyclase activating polypeptide (PACAP) References: 1(Lloyd, Maxwell et al. 1994), 2 (Deavall, Raychowdhury et al. 2000), 3(Li, Grinevich et al. 1996), 4(Voisin, Rouet-Benzineb et al. 2003), 5(May and Braas 1995), 6(Hermansen 1984), 7(Hermansen 1984), 8(Fu, Acuna-Goycolea et al. 2004), 9(Alberti, Christensen et al. 1973), 10(Björkqvist, Bernsand et al. 2005), 11 (Nowak, Maćkowiak et al. 1999), 12(Wang and Leibowitz 1997), 13(Moltz and McDonald 1985), 14(Ahren and Lundquist 1981), 15(Filipsson, Sundler et al. 1999), 16(Ahren and Lundquist 1981).

Japanese flounder *Paralichthys olivaceus* is a typical carnivorous fish species. The aim of the present study is to preliminarily investigate the brain-gut connection, and the roles of insulin and the related hormones in regulation of the glucose homeostasis in Japanese flounder loaded glucose by oral administration and intraperitoneal injection.

## 2. Materials and Methods

### 2.1. Experimental animals

Japanese flounders (body weight: 225±50 g) were provided by a local fish farm in Haiyang, Shangdong Province, China. They were randomly distributed into cylindrical fiberglass tanks (300-L) in a re-circulating water system. During the 4-week acclimation, Japanese flounders were fed with commercial feeds (Qingdao Great Bio-tech Co., Ltd) to satiation twice daily. All animal care and handling procedures were approved by the Animal Care Committee of Ocean University of China.

### 2.2. Intraperitoneal injection

There were two intraperitoneal injection (IP) groups. The one was IP injected with phosphate buffer solution (PBS), and the other one was with glucose. There were eight tanks per injection group, and eight fish per tank. Before injection, Japanese flounders were fasted for 48 hours to eliminate the effect of residual food in the digestive tract, and then were anaesthetized with MS-222 (50mg l^−1^). After being weighed, fish were IP injected with 2 ml glucose/kg body weight. The glucose was purchased from Sigma (500mg/ml, Sigma, USA). Meanwhile, the other group of animal was injected with the same volume PBS (0.01mol/L) as administration glucose group. The operations of injection for each tank were completed within 5 minutes. After injection, Japanese flounders were put back into the tanks. Samples of blood, brain, intestinal and liver were collected just before (time 0h) and 1 h, 3 h, 5 h, 7 h, 9 h, 12 h, 24 h and 48 h after injection. Blood was collected by 1ml syringes from caudal vein. Serum was obtained after centrifugation (10,000g for 10 min at 4 °C) of blood that allow to stay overnight at 4 °C. All samples were frozen in liquid nitrogen immediately and stored at -80°C.

### 2.3. Oral administration

There were two oral administration groups, and eight tanks per group, eight fish per tank. The one group received glucose (500mg/ml, Sigma, USA) with 3.34ml/Kg body mass. The other group received the same volume of PBS (0.01mol/L) and was used as the control. After oral (OR) administration, Japanese flounders were put back into the tanks. Samples of blood, brain, intestinal and liver were collected just before (time 0h), and 1 h, 3 h, 5 h, 7 h, 9 h, 12 h, 24 h and 48 h after OR. These samples were collected and stored as described above.

### 2.4. Analysis of glucose and insulin concentrations in serum

Concentration of glucose in serum was determined with the automatic biochemical analyzer (Hitachi, 7600-210, Japan). Concentration of insulin in serum was analyzed with commercial kits (Nanjing Jiancheng Bioengineering Institute, Nanjing, China).

### 2.5. Analysis of glycogen contents

The glycogen concentrations in liver were determined using the anthrone chromogenic method with commercial kits (Nanjing Jiancheng Bioengineering Institute, Nanjing, China). They were measured with the UV spectrophotometer (UV-2401PC, Shimadzu, Kyoto, Japan).

### 2.6. Tissue distribution of the selected brain-gut peptides

Tissue distributions of the selected brain gut peptides were detected in stomach, gill, spleen, muscle, brain, liver, kidney, intestine and eye by semi-quantitative RT-PCR. Specific primers for peptides and β-actin (reference gene) were shown in Table 1. Isolation of the total RNA and synthesis of the first strand cDNA were carried out following the instructions of the kit (Trizol and PrimeScript Reverse Transcriptase, Takara, Japan). The PCR reaction system consisted of 1μl of cDNA, 12.5μl 2× EsTaqMsterMix (CWbiotech, China), 9.5μl of EDPC water and 1μl of each primer (10μM). The PCR amplification program is pre-denatured at 94 °C for 5 min, followed by 30 cycles of denaturation at 94 μC for 30s, annealing at 58 μC for 30 s, extension at 72 μC for 1 min 30 s and post-extension at 72 μC for 5 min. PCR products were selected by 1.2% agarose gel electrophoresis.

**Table 1.** Expression levels of cholecystokinin precursor (Pro-CCK), neuropeptide Y (NPY), preprovasoactive intestinal peptide (Pro-VIP) and pituitary adenylate cyclase activating polypeptide (PACAP) mRNA in Japanese flounder.

|  | Pro-CCK | NPY | Pro-VIP | PACAP |
| --- | --- | --- | --- | --- |
| Eye | 0.06±0.02<sup>a</sup> | 24.42±0.33<sup>e</sup> | 1.01±0.19<sup>b</sup> | 1.04±0.39<sup>b</sup> |
| Stomach | 1.08±0.33<sup>b</sup> | 0.32±0.03<sup>ab</sup> | 4.19±0.61<sup>c</sup> | 0.02±0.00<sup>a</sup> |
| Gill | 0.01±0.00<sup>a</sup> | 4.87±1.72<sup>cd</sup> | 1.19±0.03<sup>b</sup> | 0.01±0.00<sup>a</sup> |
| Spleen | 0.00±0.00<sup>a</sup> | 5.90±1.20<sup>d</sup> | 1.03±0.01<sup>b</sup> | 0.01±0.00<sup>a</sup> |
| Muscle | 0.00±0.00<sup>a</sup> | 1.13±0.08<sup>ab</sup> | 0.02±0.01<sup>a</sup> | 0.00±0.00<sup>a</sup> |
| Brain | 24.17±5.65<sup>d</sup> | 36.82±3.32<sup>f</sup> | 8.38±1.01<sup>c</sup> | 8.47±0.77<sup>c</sup> |
| Liver | 0.03±0.02<sup>a</sup> | 1.54±1.02<sup>ab</sup> | 0.22±0.24<sup>a</sup> | 0.01±0.01<sup>a</sup> |
| Kidney | 0.00±0.00<sup>a</sup> | 0.04±0.01<sup>a</sup> | 0.03±0.01<sup>a</sup> | 0.00±0.00<sup>a</sup> |
| Intestine | 5.01±0.69<sup>c</sup> | 2.18±0.72<sup>bc</sup> | 14.54±1.44<sup>d</sup> | 17.81±2.77<sup>d</sup> |

### 2.7. Gene expression analysis by real-time quantitative RT-PCR

Gene expression levels were determined by real-time quantitative RT-PCR (q-PCR) using the iCycler iQ™ (Bio-Rad,Hercules, CA, USA). Analyses were using SYBR Green I (CWbiotech, China) according to the manufacturer’s instructions. The total reaction volume was 25μl, including 1.0μl cDNA, 12.5μl 2× UltraSYBR Mixture, 1μl each of gene-specific primer (10μM, Table2) and 9.5μl DEPC water. Thermal cycling was initiated with incubation at 95 °C for 10min using hot-start iTaq™ DNA polymerase activation, 40steps of PCR were performed, each one consisting of heating at 95 °C, 10s for denaturing, and at specific annealing and extension temperatures is Tm for 30s, 72°C for 32s. Following the final PCR cycle, melting curves were systematically monitored (58 °C temperature gradient at 0.5 °C/s from 58 to 95 °C) to ensure that only one fragment was amplified. Relative quantification of the target gene transcript was done using β-actin gene expression as reference (Olsvik, Lie et al. 2005) which was stably expressed in this experiment. Samples without reverse transcriptase and samples without RNA were run for each reaction as negative controls. Relative quantification of the genes were calculated using “2^-ΔΔCt^” meth (Livak and Schmittgen 2001) with β-actin as reference gene. All real-time Q-PCR was performed in triplicate biological replicates.

**Table 2.**
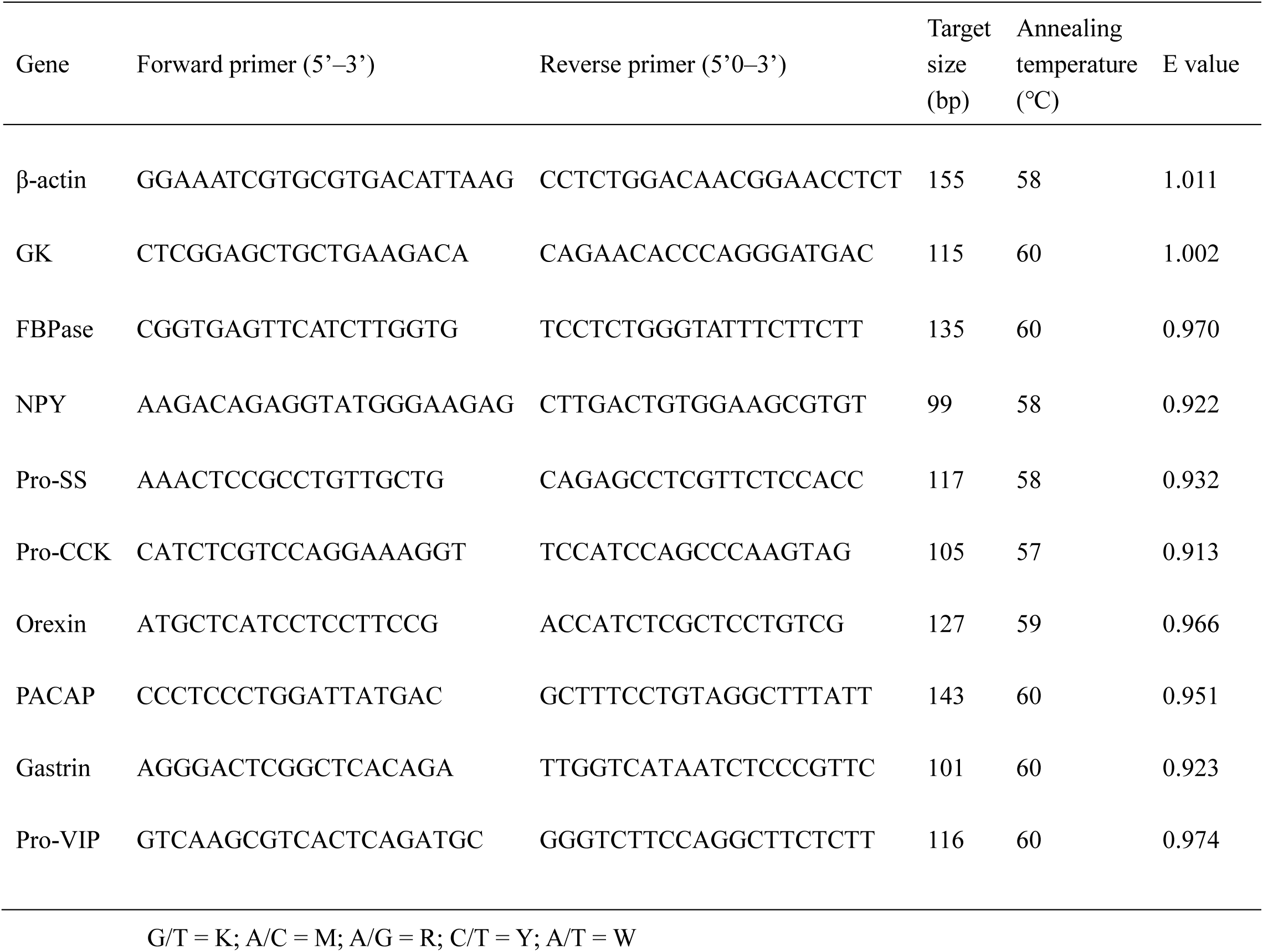
Primers used for gene mRNA quantification by RT-PCR.

The effects of glucose administration on the expression of hormone (peptides) genes were detected by comparing the 0, 1, 3, 5, 7, and 24 h after intraperitoneal injection and oral glucose administration, respectively. Hepatic glucose metabolism-related genes were analyzed by comparing the 0 h, 5h and 24 h in the treatment of intraperitoneal injection of glucose, and 0 h, 3 h and 12 h in the oral glucose administration.

### 2.8. Statistics analysis

All data were expressed as means ± standard error and performed using SPSS 17.0. A one-way analysis of variance (ANOVA) was used to compare the differences in relative brain-gut peptides gene expression over time. When overall differences were considered statistically significant at *P* < 0.05, Tukey’s test was used to compare the means among individual treatments.

## 3. Results

### 3.1. Concentrations of glucose and insulin in serum

Concentrations of glucose in serum reached the peak (20.06±1.92mM) at 5h after the IP injection of glucose, which was about 28 times as high as that at 0h (0.71±0.25mM) (Figure 2A). From the 5h to 24h after injection, the blood glucose decreased to 7.40±5.47mM. There was no significant difference in blood glucose between the time point of 0h and 24h (Figure 2A). After injection of glucose, the insulin concentration in serum was decreased to the lowest value (1.58±0.21mIU/L) at 3h (Figure 2B). After that, it grew gradually to the normal value at 24h (3.03±0.006mIU/L) as that at 0h (3.20±0.18 mIU/L).

**Figure 2.**
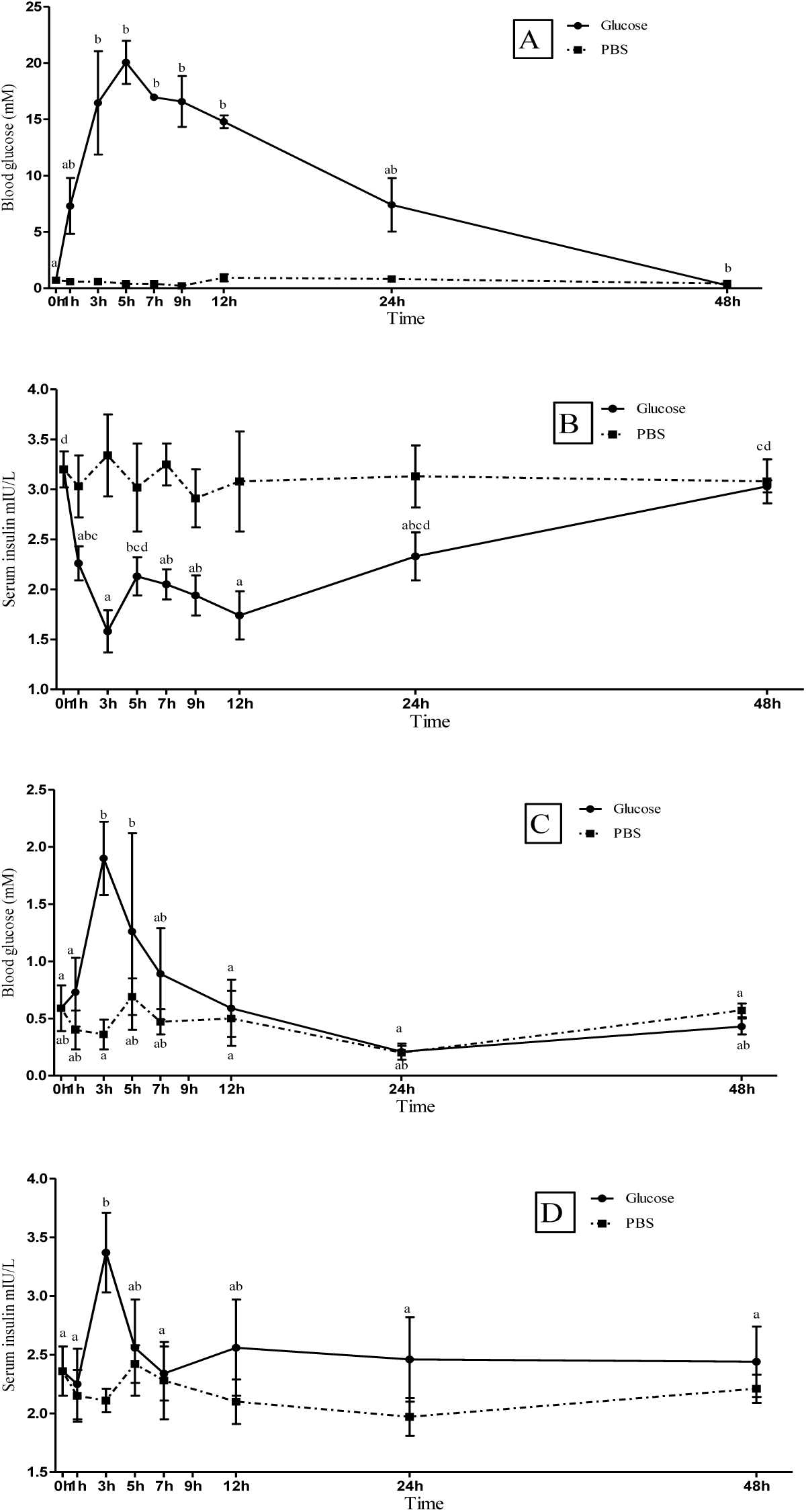
The concentration of blood glucose and serum insulin in Japanese flounder after intraperitoneal (IP) (A, B) and oral (OR) (C, D) administrationof glucose or phosphate buffer solution (PBS). Each value is expressed as the Mean ± SE (n=3). Values sharing a common superscript letter were not significantly different.

After oral administration of glucose, blood glucose concentration had rapidly increased at 3h (1.90±0.23mM), and it was higher than those at the other time points. After the 3^rd^ hour, the blood glucose concentration decreased gradually (Figure 2C). The serum insulin concentration increased from 2.36±0.21mIU/L (0h) to 3.37±0.34mIU/L (3h). It returned to the normal level after 5h, and then, kept relatively stable (Figure 2D).

### 3.2. Tissue distribution of peptides genes

Results of the gene distribution and expression are shown in Figure 3 and Table 1. The relative expressions of prosomatostatin (Pro-SS) and orexin precursor (Pro-OX) in brain were relatively higher than that in the other analyzed tissues including stomach, gill, spleen, muscle, liver, kidney, intestine and eyes. Gastrin was mainly expressed in intestine (Figure 3). NPY, CCK precursor (Pro-CCK), Preprovasoactive intestinal peptide (Pro-VIP) and PACAP were detected in many tissues (Figure 3). Relative expression of NPY had the highest value in brain, and followed by the eye, gill and spleen. Relative expression of Pro-CCK was up to maximum in brain, followed by the intestine and stomach. The expression of Pro-VIP was relatively high in intestine, but low in brain, stomach, gill, spleen and eyes. The PACAP had the highest expression in intestine, then brain.

**Figure 3.**
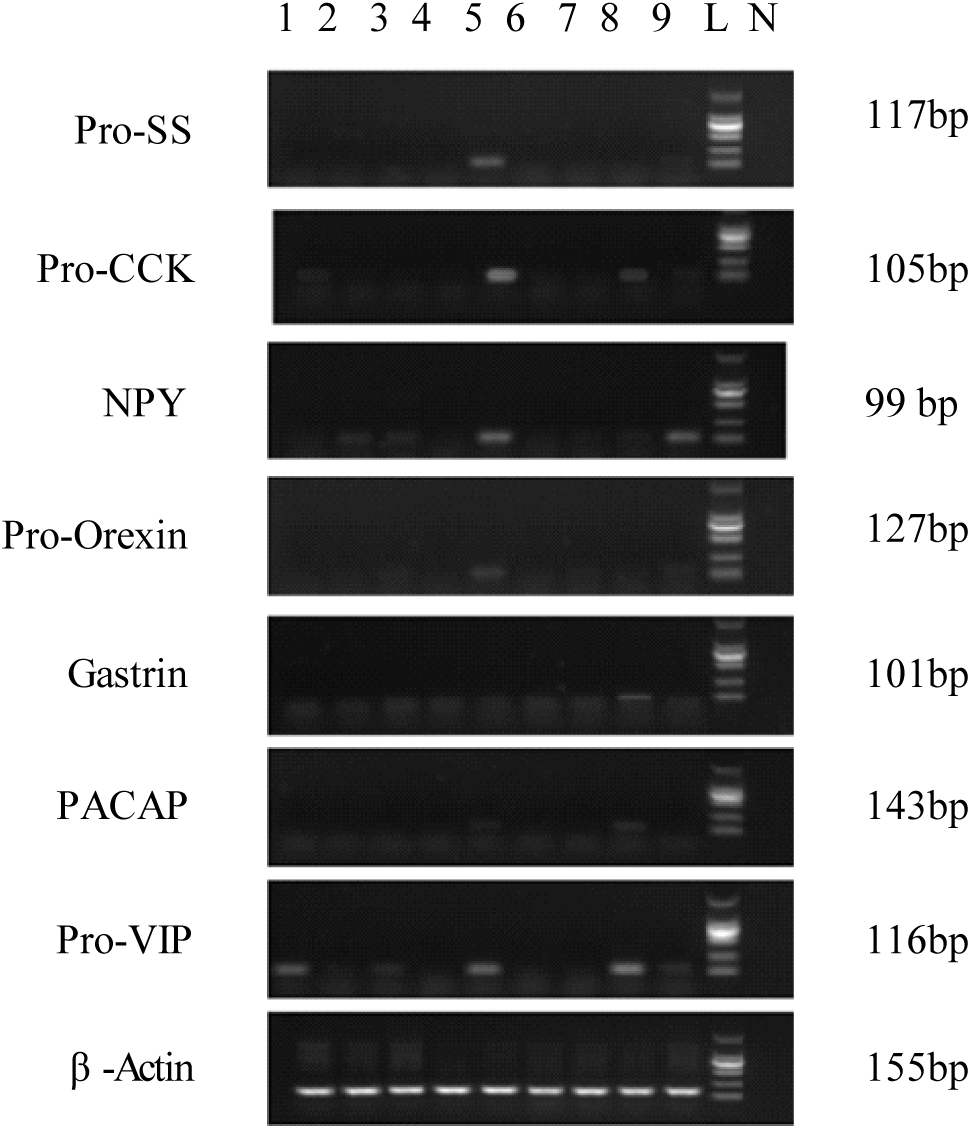
Tissue distributions of brain-gut peptides genes. L: ladder, 1: stomach, 2: gill, 3: spleen, 4: muscle, 5: brain, 6: liver, 7: kidney, 8: intestine, 9: eyes, N: negative control. Expression of housekeeping gene β-actin was observed to ensure the integrity of the cDNA template of each tissue sample. Prosomatostatin (Pro-SS), cholecystokinin precursor (Pro-CCK), neuropeptide Y (NPY), orexin precursor (Pro-OX), pituitary adenylate cyclase activating polypeptide (PACAP), preprovasoactive intestinal peptide (Pro-VIP)

### 3.3. Gene expressions after glucose administration

#### 3.3.1. Gene expression of the Pro-SS, NPY, Pro-CCK and Pro-OX in brain

Results of the gene expression in brain after glucose administration are shown in Figure 4.

**Figure 4.**
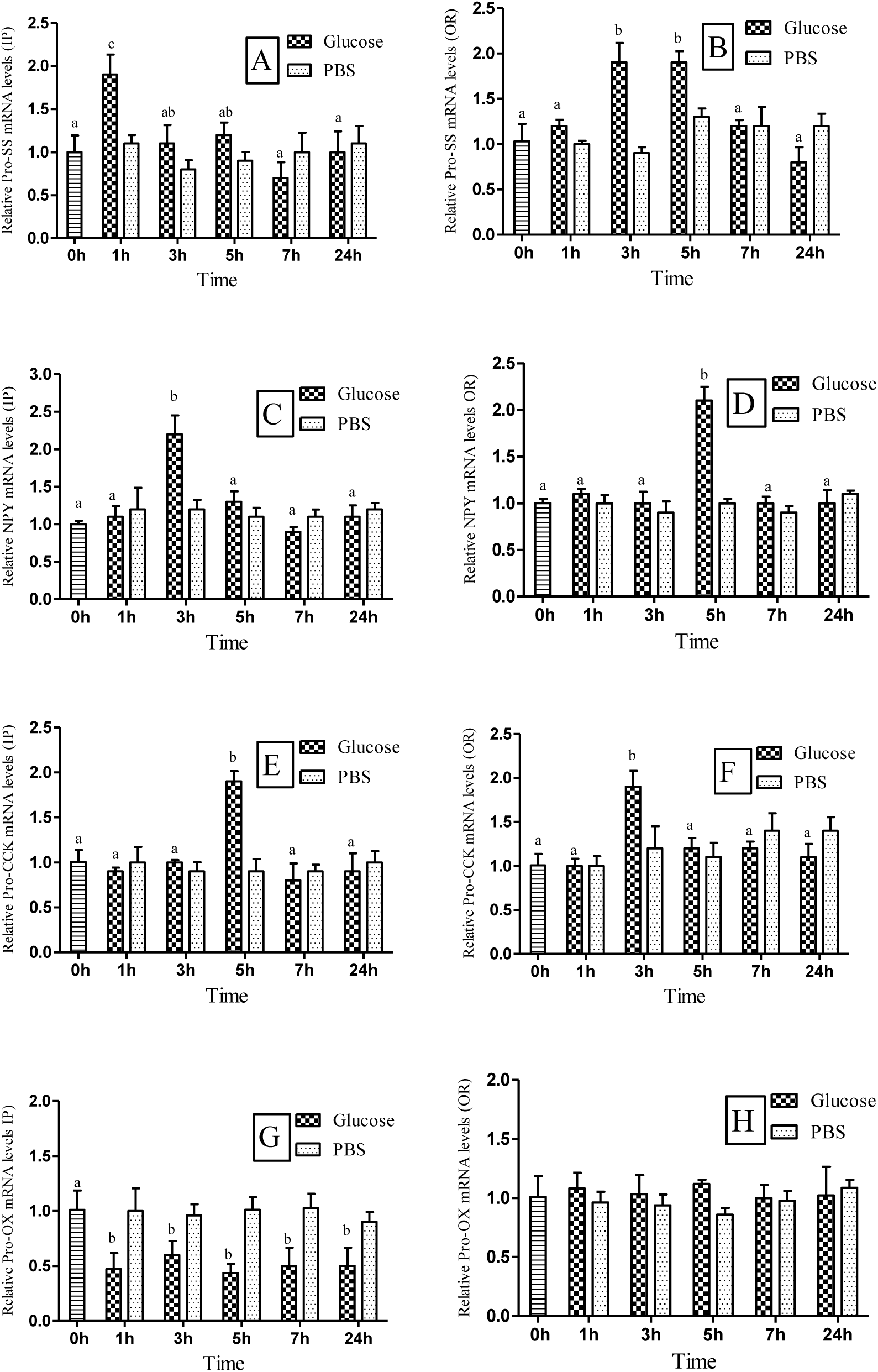
Gene expressions of prosomatostatin (Pro-SS), neuropeptide Y (NPY), cholecystokinin precursor (Pro-CCK) and orexin precursor (Pro-OX) in the brain of Japanese Flounder after intraperitoneal (IP) (A, C, E, G) and oral (OR) (B, D, F, H) administration of glucose (Glu) or phosphate buffer solution (PBS). Each value is expressed as the Mean ± SE (n=3). Values sharing a common superscript letter were not significantly different.

One hour after the intraperitoneal injection of glucose, mRNA levels of the Pro-SS were significantly increased, then it was not significantly differed from that at 0h at the others time points. In the group of IP PBS, there were no significant different in mRNA levels of Pro-SS among all the time points (Fig.4A).

After oral glucose administration, Pro-SS mRNA levels were significantly up-regulated at the time point of 3h and 5h (Fig.4B). The NPY mRNA levels were significantly increased only at 3h after IP and 5h after OR glucose (Fig.4C, 4D). The cholecystokinin precursor (Pro-CCK) mRNA levels were significantly increased only at 5h after IP and 3h after OR glucose (Fig. 4E, 4F). The Pro-OX mRNA levels significantly decreased at all the time points after IP glucose. However, there were no significant differences among all the time points after OR glucose (Fig.4G, 4H).

#### 3.3.2. Gene expression of Pro-VIP, PACAP and gastrin in intestine

Results of the gene expression in intestine after glucose administration are shown in Figure 5.

**Figure 5.**
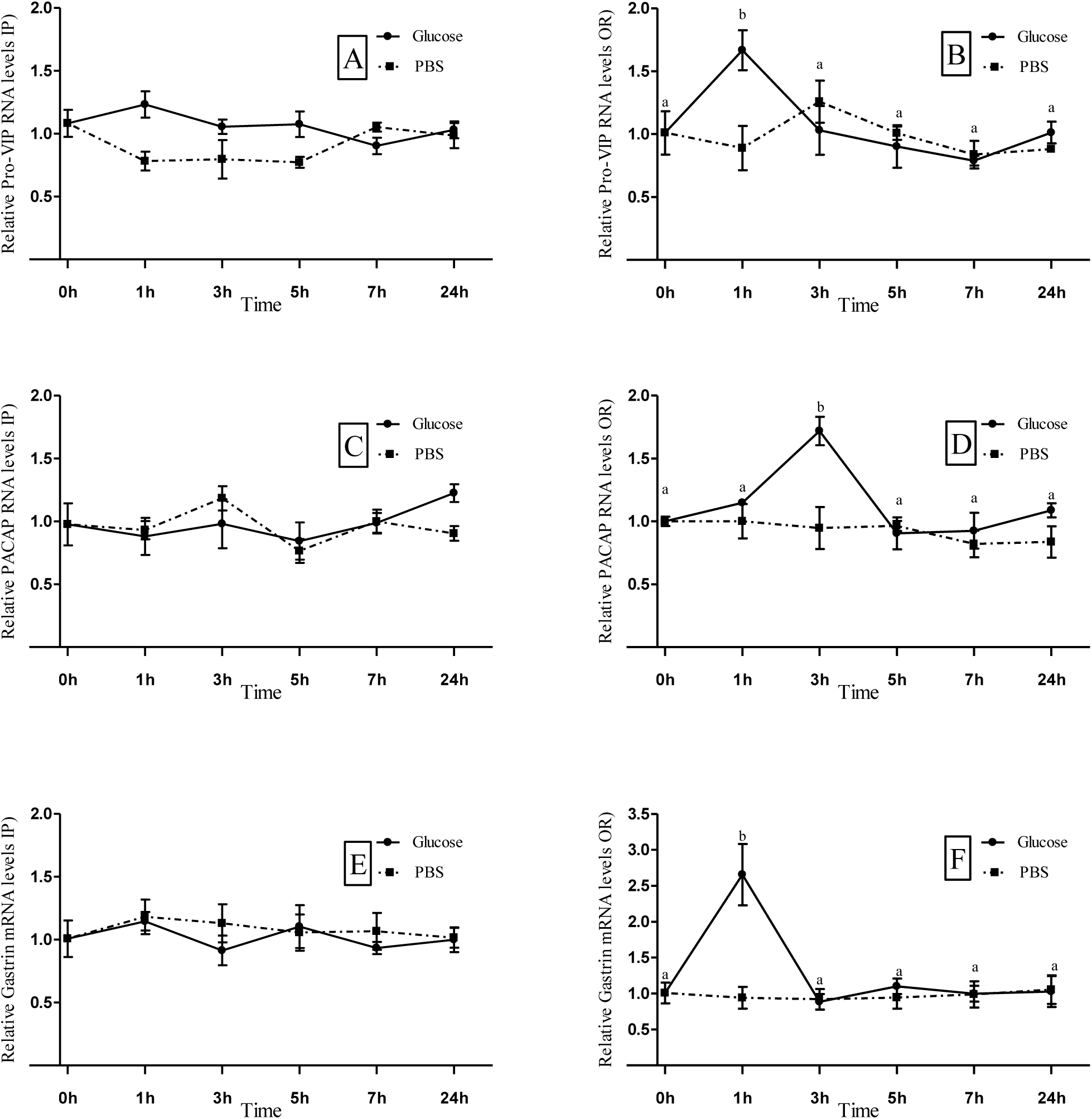
The gene expression of preprovasoactive intestinal peptide (Pro-VIP), pituitary adenylate cyclase activating polypeptide (PACAP) and gastrin in the gut of Japanese Flounder after intraperitoneal (IP) (A, C, E) and oral (OR) (B, D, F) administration of glucose (Glu) or phosphate buffer solution (PBS). Each value was expressed as the Mean ± SE (n=3). Values sharing a common superscript letter were not significantly different.

There were no significant differences in gene expressions of preprovasoactive intestinal peptide (Pro-VIP), PACAP and gastrin among all the time points after IP glucose.

There was only one time point after OR glucose at which that the expressions of these three genes were significantly higher than that at 0h. The time point was 1h, 3h and 1h, respectively. There were no significant differences between the rest time points and 0h.

### 3.4. Heatmap of glucose affected brain-gut peptides gene expression

The expression of brain-gut peptides gene in brain and intestines of Japanese flounder by different ways of glucose administration was analyzed on heatmap (Fig.6). As shown in Figure 6, it consists of two treatment groups: intraperitoneal injection glucose (1g/Kg) and oral glucose administration (1.67/Kg). The results show that in intraperitoneal injection glucose (1g/Kg), the gene expressions of Pro-SS, NPY, Pro-CCK were increased at 1h, 3h, 5h compared to 0h respectively, and Pro-OX expression was declined after 1h. In oral glucose group (1.67g/Kg), Pro-SS, NPY, Pro-CCK expression increased at 3-5h, 5h and 3h respectively, and the mRNA levels of Pro-VIP, PACAP, Gastrin were promoted at 1h, 3h and 1h, respectively.

**Figure 6.**
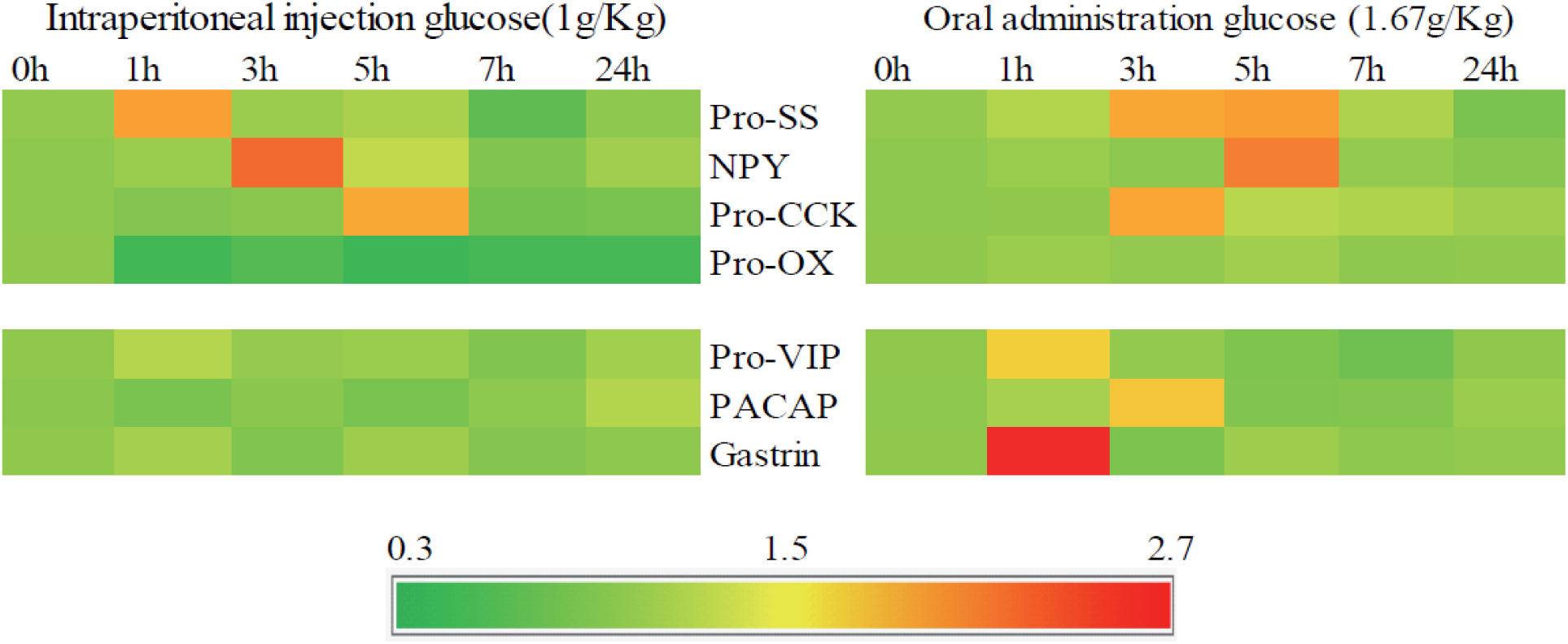
Visualizing gene expression of prosomatostatin (Pro-SS), neuropeptide Y (NPY), cholecystokinin precursor (Pro-CCK), orexin precursor (Pro-OX), preprovasoactive intestinal peptide (Pro-VIP), pituitary adenylate cyclase activating polypeptide (PACAP) and gastrin on heatmap Note: It consists of two treatment groups: intraperitoneal injection glucose (1g/Kg) and oral glucose administration (1.67/Kg). In this figure, the graphical presentation of data, numerical values are displayed by colors.

### 3.5. Gene expression of glucokinase (GK) and fructose-1,6-bisphosphatase (FBPase) in liver

Results of the gene expression of GK and FBPase in liver after glucose administration are shown in Figure 7. The fish showed the highest GK mRNA level after IP and OR glucose at 5h and 3h, respectively (Fig.7A, 7B). The FBPase mRNA level significantly decreased at 5h and 3h after IP and OR glucose, respectively. And then increased at 24h and 12h, respectively, and had no significant differences with that at 0h.

**Figure 7.**
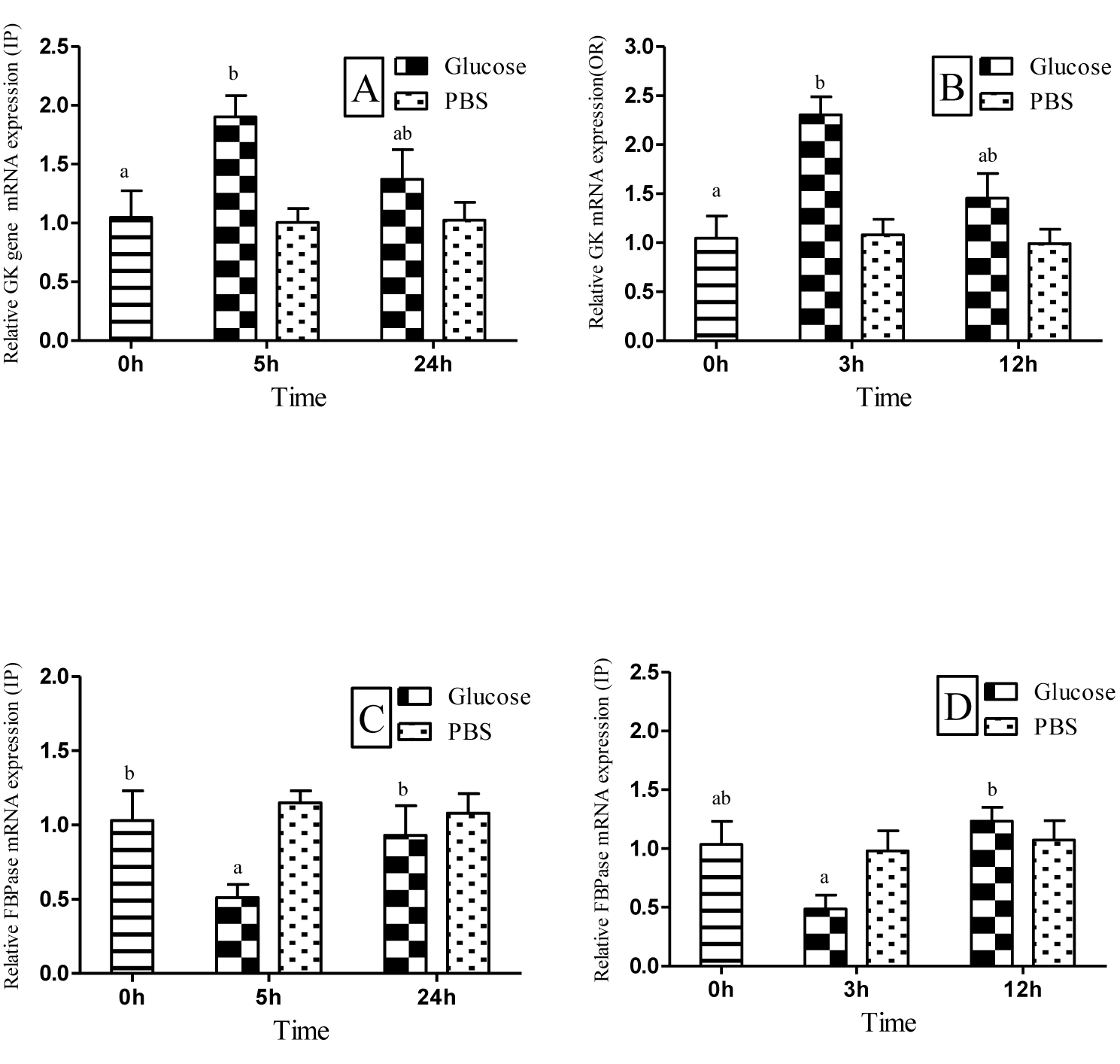
The gene expression of glucokinase (GK) and fructose-1,6-bisphosphatase (FBPase) in the liver of Japanese Flounder after intraperitoneal (IP) (A, C) and oral (OR) (B, D) administration of glucose (Glu) or phosphate buffer solution (PBS). Each value was expressed as the Mean ± SE (n=3). Values sharing a common superscript letter were not significantly different.

### 3.6. Glycogen in liver

The changes of liver glycogen contents are shown in Figure8. In the IP glucose group, liver glycogen content decreased at 3h and increased after 5h. And the significant highest value was found at 9h. In the OR glucose group, the highest value of liver glycogen content was found at 5 h. By contrast, the liver glycogen contents in the PBS groups showed no significant differences among all the time points.

**Figure 8.**
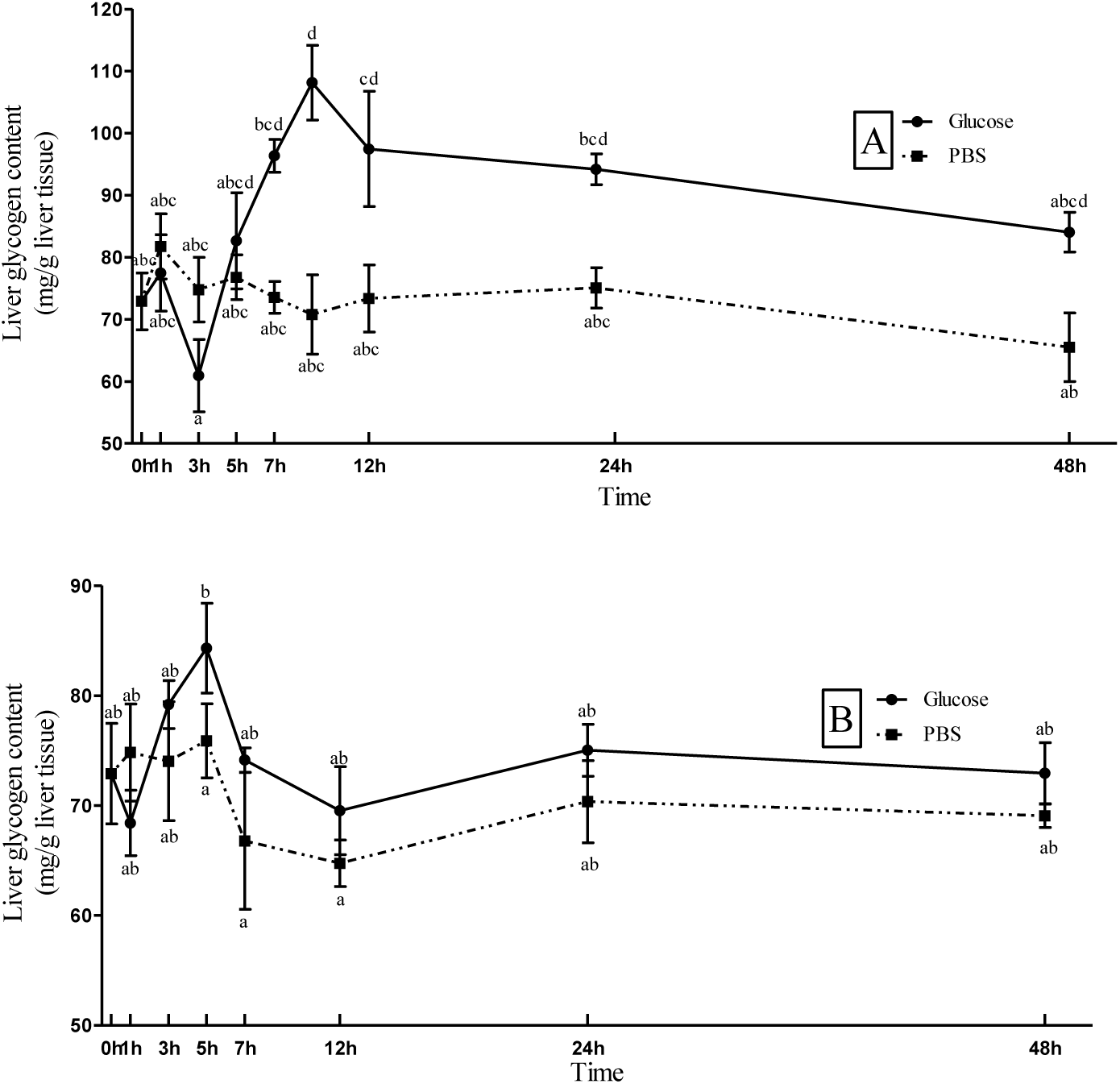
The glycogen contents in liver of Japanese flounder after intraperitoneal (IP) (A) and oral (OR) (B) administration of glucose or phosphate buffer solution (PBS). Each value was expressed as the Mean ± SE (n=3). Values sharing a common superscript letter were not significantly different.

## 4. Discussion

### 4.1 Blood glucose levels changed by different ways of glucose administration

The present study showed that the blood glucose content of Japanese flounder reached its peak (20.06±1.92mM) at 5h after intraperitoneal injection of glucose (500mg/ml, 1g/Kg) and hyperglycemia lasted about 21 hours. This was similar with the previous study on Australian snapper *Pagrus auratus*, in which the blood glucose reached to the peak at 3h (18.9mM) and the hyperglycemia lasted for about 18 hours (Booth, Anderson et al. 2006). However, after intraperitoneal injected same or even a higher dose of glucose, some omnivorous fish like tilapia, white sea bream, spends from 1-2 hours reaching to the peak in the concentration of blood glucose and 6-9 hours recovering as usual (Wright Jr, O’Hali et al. 1998, Enes, Peres et al. 2012). It was suggested that the omnivorous fish species had higher ability of blood glucose control than the carnivorous. After oral administration of glucose (500mg/ml, 1.67g/kg), blood glucose of Japanese flounder peaked (1.90±0.23mM) at 3h, and it returned to normal level (0.89±0.04mM) at 7h. Duration of hyperglycemia is about 5 hours. These results are similar to those previous researches in carnivorous grouper (1.67g/Kg) (Yang, Ye et al. 2012). As carnivorous fish, however, the duration of hyperglycemia is about 10 hours and 36 hours in black carp *Mylopharyngodon piceus* and Chinook salmon *Oncorhynchus tshawytscha* after oral administration of glucose (1.67g/Kg), respectively. It was suggested that in carnivorous fishes, glucose tolerance significantly differed in species. In addition to the species difference, the culture environment and conditions could also be part of the reasons. As an omnivorous fish, after oral administration of glucose (1.67g/Kg) in allognogenetic gibel carp, the blood glucose concentration reached the peak value (26.04mM) at 3h, and the hyperglycemia lasted about 7 hours (Ying 2003). As a herbivorous fish, after oral administration of glucose (1.67g/Kg) in grass carp *Ctenopharyngodon idellus*, the highest blood glucose concentration reached at 3h, and the hyperglycemia lasted about 6 hours (Huang, Ding et al. 2005). It was suggested that the carnivorous fish had lower ability of blood glucose control than the omnivorous and herbivorous.

In the present study, the hyperglycemia lasted 5 hours in oral administration of glucose group, and the highest concentration of blood glucose was 1.90±0.32mM. While in the IP injection of glucose group, the hyperglycemia lasted 21 hours, and the peak value of blood glucose was 20.06±2.72mM. It was suggested that the Japanese flounder has stronger capacity in eliminating glycaemia caused by oral administration of glucose than by the IP injection of glucose. This could be partly due to the following reasons. In OR glucose group, a portion of glucose could be consumed when glucose enters gastrointestinal tract. While the left portion of glucose was absorbed into bloodstream. If high glucose loads in this experiment caused pathology, oral glucose may likely discharge to the vitro through digestive tract. Glucose by the way of intraperitoneal injection was mostly absorbed into the bloodstream directly by the peritoneal capillary.

In IP injection glucose, the content of insulin declined as blood glucose rose (3h-24h), from 0h to 3h. After the third hour, the insulin did not recover at the original value at all. However, in oral glucose administration, as the blood glucose going up (1h-7h), the insulin was growing as well (1h-5h). Therefore, the different tendency of insulin is also the sound reason of short last of hyperglycemia in oral glucose administration.

### 4.2 Serum insulin changed by different ways of glucose administration

In the present study, serum insulin level sharply declined in intraperitoneal injection of glucose group, and then increased. The results are similar to the previous study in rainbow trout with IP injection of glucose (Harmon, Eilertson et al. 1991) and grouper *Epinphelus Coioides* by OR administration of glucose (Yang, Ye et al. 2012).

In Japanese flounder, IP glucose was mostly absorbed in the bloodstream through the peritoneal capillary. This leads to the continuous rise of blood glucose. Glucose comes to the brain through blood circulation. Due to the arcuate nucleus in the hypothalamus of the blood-brain barrier permeability being high, after the stimulating of glycemia to brain, the gene expression of Pro-SS, NPY, Pro-CCK and Pro-OX was significantly different at 1h, 3h, 5h and 1h compared to 0h, respectively. It was suggested that the sensitivity of these hormones in the brain was different. This also shows that in the period of blood glucose raised by IP injection glucose, the hormones play different roles. Combining the function of these hormones in mammal, it was suggested that insulin declined at 3h after IP injection glucose administration could be relevant to these hormones.

The gene transcript of Pro-SS and NPY was up-regulated at 1h and 3h, respectively after IP glucose. The function of the SS and NPY is to inhibit the secretion of insulin. Pro-OX expression was down-regulated at 1h. The function of OX is to enhance insulin production which is stimulated by glucose (Nowak, Maćkowiak et al. 1999). That is to say the secretion of insulin was depressed in the first 3 hours after IP glucose administration. After 3h of IP glucose, insulin levels rose. Simultaneously, the gene expression of Pro-CCK significantly elevated. The CCK function is to stimulate insulin secretion (Hermansen 1984). After intraperitoneal injection, glucose didn’t enter into intestinal but directly into abdominal cavity capillary, so there was no significant difference in expression of gut hormone genes, such as Pro-VIP, PACAP and gastrin (Fig 9A).

**Figure 9.**
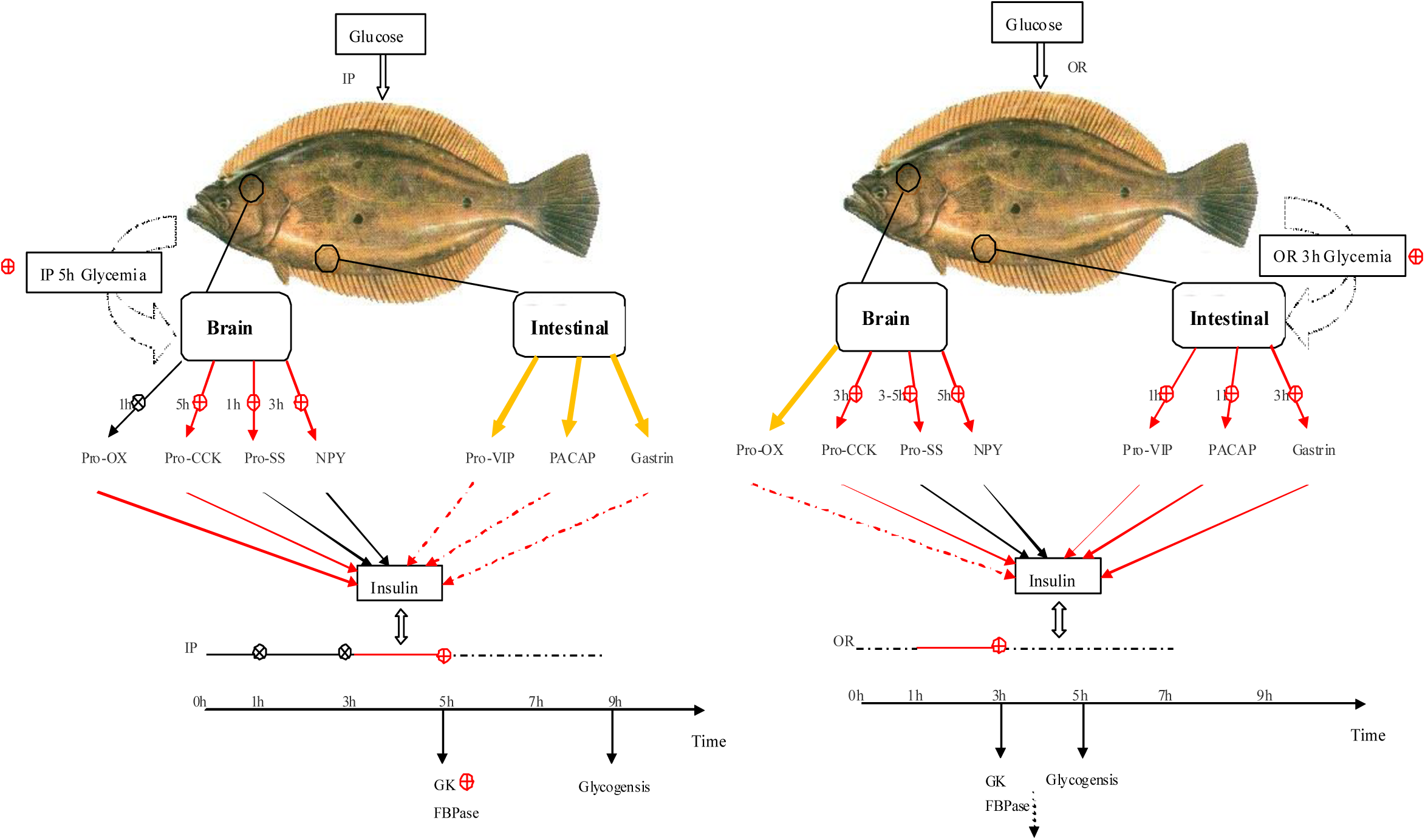
Summary on the results of the present study. Regulations of glucose metabolism in brain and intestine were affected by intraperitoneal (IP) (A) and oral (OR) (B) administration of glucose. Red line stands for stimulation, and black line means inhibition. The red circle represents the significant promotion effects. The black circle represents the significant inhibition effects. Black dotted line means inhibition, but did not show significiant difference. The yellow line stands for there is no significant difference in all time points. Point in time after glucose loads with red and black symbol means significant difference (*P*<0.05) was found at that time point.

In OR glucose group, glucose enters into gastrointestinal tract, while parts of glucose were consumed by cells and other parts of glucose were absorbed into the bloodstream. After oral glucose administration, insulin increased at 1-3h, meanwhile it begun to drop after 3h. Pro-CCK expression which could promote the secretion of insulin runs up at 3h. The differences of expression quantity of Pro-SS occurred at 3-5h, while for NPY the difference appeared at 5h. Both SS and NPY can inhibit the secretion of insulin in mammals. Before high glycemia appears, hormones in gut which can stimulate insulin secretion like VIP (Ahren and Lundquist 1981), PACAP (Filipsson, Sundler et al. 1999), gastrin (Ahren and Lundquist 1981) were high in expression at 1h, 1h and 3h, respectively. In mammals, when food enters into intestine, intestinal mucosal cells can secrete a variety of gastrointestinal hormones, which have an important effect on the secretion of insulin. These hormones include glucose-dependent insulinotropic polypeptides, glucagon-like peptide-1, gastrin, etc. As the intestinal signals, it can stimulate insulin secretion before glycemia appears to align the body absorbed nutrient.

In mammals, SS, NPY, CCK, OX, VIP, PACAP and gastrin belong to the brain-gut peptide hormones, and they can be found in the brain and intestines. In the present study, only the Pro-CCK, Pro-VIP and PACAP were found both in the brain and intestine. The expressions of Pro-SS and Pro-OX were only detected in the brain. And the expression of gastrin was only found in the gut. This could be due to the less secretion of these hormones. Anyway, it was confirmed that the brain-gut peptides and their gene expression were detected in brain and gut. This could support the hypothesis of brain-intestine connection in Japanese flounder. In mammals, the function of CCK in the gut is to control the release of pancreatic enzyme and gallbladder contraction. It acts as a neurotransmitter, which can control feeding, analgesia, blood pressure, memory, insulin release in the nervous system (Du Jing 2007). Pro-VIP and PACAP are mainly as neurotransmitter in brain. In the digestive system, they mainly act as gastrointestinal hormones, and promote insulin secretion (Filipsson, Sundler et al. 1999, Xu Wei 2002, Zhang Yang 2009).

The reason could be that in the early stage of glucose administration, the blood glucose increased sharply. It could cost large number of insulin original in the body. At the same time, the insulin secretion stimulating by glucose administration could be less than the costing. So the serum insulin concentration declined sharply.

The insulin content in group of IP glucose as a whole tends to increase after 3h, but it does not exceed value at 0h. The result, with the value higher than that at 0h, is different from that in OR group. This may in that the apoptosis of β-cell leads to the decrease of the function of insulin

In my experiments, serum insulin will increase with blood glucose rise. Therefore, we can get a conclusion that insulin changes are the result of a combination of blood glucose and brain-gut peptides.

### 4.3 Metabolism responses of the glycolysis and gluconeogenesis in liver

Glycolysis and gluconeogenesis mainly participate in the catabolism and synthesis of carbohydrate, coordinating with each other to ensure the glucose homeostasis of fish (Nie Q, 2013). As one of the hexokinase (HK) isoenzymes, GK is the initial enzyme of the glycolytic pathway, and the FBPase is one of the key enzymes in the gluconeogenesis pathway. In the present study, the gene expression of GK was significantly enhanced both after IP injection and OR glucose. Similar results were found in turbot (Nie, Miao et al. 2015), gilthead sea bream (Metón, Caseras et al. 2004) and common carp (Panserat, Rideau et al. 2014). Compared to the GK, the gene expression of FBPase was significantly decreased in the treatment of intraperitoneal injection at 5 h. Meanwhile, the similar results were found in the group of oral glucose. Researches in European sea bass found that dietary starch (10% to 30%) could not affect the FBPase activity and gene expression (Moreira, Peres et al. 2008). In omnivores like common carp, Panserat, Plagnes-Juan et al. (2002) also reported that there are no significant differences in FBPase gene expression between the fish fed with carbohydrates or not. The result suggested that the gene expression was irrespective to the carbohydrate intake. Furthermore, (Fernández, Miquel et al. 2007) it also found that dietary cornstarch (5% to 26%) could not affect the FBPase gene expression in gilthead sea bream, whereas Panserat, Plagnes-Juan et al. (2002) found that the gene expression of FBPase significantly decreased after fed dietary carbohydrate (20%).

Researches in rainbow trout, tilapia, European sea bass and gilthead sea bream showed that the liver glycogen content significantly increased when fed dietary carbohydrates (Mazur, Higgs et al. 1992, Shiau and Liang 1995, Couto, Enes et al. 2008, Moreira, Peres et al. 2008). In the present study, the reduction in liver glycogen content during the first 3 hours after injection or 1 hour after oral glucose might be explained by the effect of glycogenolysis in fish. To some degree, this was similar to the previous finding in turbot (Garcia-Riera and Hemer 1996). Peres, Goncalves et al. (1999) found that liver glycogen content of seabream significantly decreased during the first hour after glucose injection, while the liver glycogen content of seabass increased significantly. In present study, IP glucose group, liver glycogen content increased after 5h. And the significant highest value was found at 9h. In the OR glucose group, the highest value of liver glycogen content was found at 5 h. The liver glycogen content reached their maximum behind the occurrence of blood glucose peak. This is an evidence for Japanese flounder to turn glucose into glycogen. It could be a strategy to ease the stress of high blood glucose in Japanese flounder.

In the present study, when the blood glucose concentration peaked in both IP and OR group, the GK was stimulated and the FBPase was depressed. This could be the responses to decrease the hyperglycemia. In the IP glucose group, liver glycogen content decreased at the 3h, then went up at the 5h, peaked at the 9h. In the OR glucose group, the highest value of liver glycogen content was found at the 5h. It is obviously that the time of glycogen synthesis and reaching peak was later than those of hyperglycemia. It is suggested that organism reduces blood glucose by glycogen synthesis.

The function of insulin in glucose metabolize is through insulin receptor substrate-1 (IRS-1), activation of PI_3_K, and consequent Akt phosphorylation, further phosphorylate downstream signaling protein, such as AS160, mTORC1, FoxO1 and GSK3. phosphorylation of AS160 is required for GLUT4 translocation and it can promote tissue uptake glucose (Sano, Kane et al. 2003). mTORC1 activation is sufficient to stimulate glycolysis, the oxidative arm of the pentose phosphate pathway, and promote the decomposition of glucose (Düvel, Yecies et al. 2010). Insulin levels increase upon feeding and signal through Akt to suppress Foxo1 by phosphorylation and exclusion from the nucleus. gluconeogenesis genes are regulated by Foxo1 (Gross, Wan et al. 2009). Akt phosphorylates and inactivates GSK-3 increased GLUT1 levels and in enhanced glucose uptake through these high-affinity transporters. (Buller, Loberg et al. 2008) Insulin signal pathway also promote glycogen synthesis by inactivates GSK-3 though phosphorylates Akt (Lochhead, Coghlan et al. 2001).

In conclusion, in the present study, brain-gut peptides were confirmed in the present study. And the serum insulin and the brain-gut peptides have different responses between the IP and OR administration of glucose. A negative feedback mechanism in the insulin-regulated glucose homeostasis was suggested in Japanese flounder. Furthermore, this regulation could be conducted by activating PI3k-Akt, and then lead to the pathway downstream changes in glycolysis and gluconeogenesis. This supposition was expressed in Figure 10.

**Figure 10.**
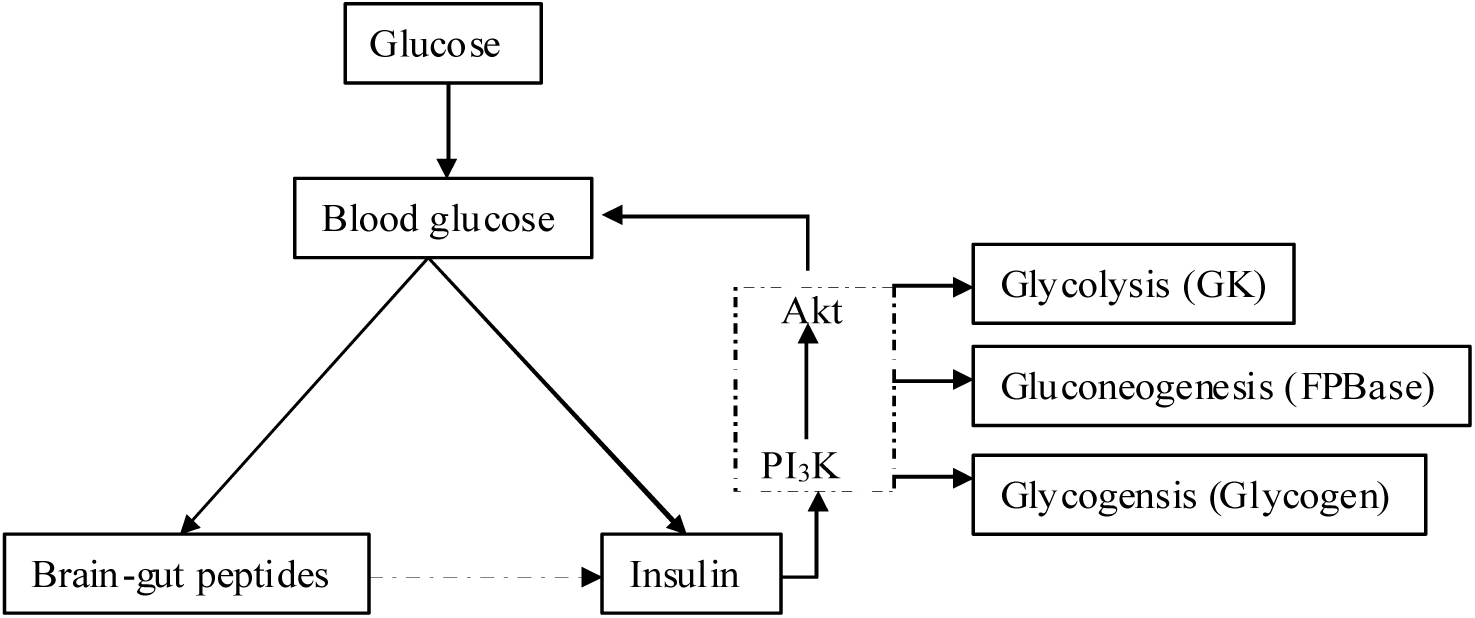
A supposed negative feedback regulation mechanism of glucose homeostasis controlled by insulin and brain-gut peptides. The insulin could decrease the blood glucose level of Japanese flounder by activating PI3k-Akt, and then it could also cause the pathway downstream changes in glycolysis, gluconeogenesis and glycogenesis. The dashed and dashed boxes represent the relationships that exist in mammals, which may exist in Japanese flounder.

## Acknowledgements

This study was financially supported by the National Basic Research Program (973 program, No. 2014CB138600), the National Natural Science Foundation of China (No. 31572628) and the Fundamental Research Funds for the Central Universities of Ocean University of China (No. 201562017).

## Author contributions

W.B.Z. and K.S.M. designed the experiment and revised the manuscript. D.L., D.D.H., B.Y.G, K.Y.D., Z.X.G., M.X.Y. and W.X. completed the experiment and analyzed the data. D.L. and D.D.H. prepared the manuscript. All authors were involved in the discussion of experimental data.

